# Examining the relationship between social cognition and neural synchrony during movies in children with and without autism

**DOI:** 10.1101/2020.03.06.981415

**Authors:** KM Lyons, RA Stevenson, AM Owen, B Stojanoski

## Abstract

Children who have been diagnosed with autism spectrum disorder (ASD) often show a marked deficit in measures of social cognition. In autistic adults, measures of social cognition have been shown to relate to differences in brain synchronization (as measured by fMRI) when individuals are processing naturalistic stimuli, such as movies. However, whether children with social impairments, with or without a diagnosis of ASD, differ in their neural responses to movies has not yet been investigated. In the current study, neural synchrony, measured using fMRI, was examined in three groups of children aged 7 to 12, who differed with respect to scores on a measure of autistic traits associated with social impairment and whether or not they had been diagnosed with ASD. While watching the movie ‘Despicable Me’, those diagnosed with ASD had significantly less neural synchrony in areas that have been previously shown to be associated with social cognition (e.g. areas related to ‘theory of mind’), and plot following (e.g. the lateral prefrontal cortex), than those who did not have an ASD diagnosis. In contrast, two groups who differed in their degree of social impairment, but did not have a diagnosis of ASD, showed no significant differences in neural synchrony across the whole brain. These results shed some light on how autistic traits may contribute to an individual’s conscious experience of the world, and how, for children with ASD, that experience may differ markedly from that of those without ASD.

## Introduction

Autism spectrum disorder (ASD)^1^ is a complex developmental condition characterised by a variety of neurological and psychological features; however, the most prominent feature of ASD is a marked deficit in ‘social cognition’. Social cognition refers to understanding what other people believe, how they will react in situations, and why they feel the way they do, and is a core element of successful human interactions. Autistic individuals perform poorly on tasks that assess social cognition, such as face perception (Spencer et al., 2011), perspective taking (Hamilton et al., 2009), and theory of mind (ToM), or the ability to attribute mental states to oneself and others (Pedreño et al., 2017). One of the most common tools to screen for deficits associated with ASD is the Social Responsiveness Scale, which measures aspects of social awareness, communication, and motivation (Constantino & Gruber, 2012).

The brains of autistic individuals often show differences when compared to those of typically-developing individuals. These include structural abnormalities (Barnea-Goraly et al., 2004; Brieber et al., 2007), functional differences during task-based fMRI (Bölte et al., 2008; Gilbert et al., 2008; Just et al., 2007; Mason et al., 2008; Solomon et al., 2009) and changes in resting-state functional connectivity (Cherkassky et al., 2006; Kana et al., 2015; Monk et al., 2009; Weng et al., 2010). Many of the brain regions that show differences in autistic individuals have been linked to ToM in healthy individuals, including the temporal parietal junction (Saxe & Kanwisher, 2003; Saxe & Wexler, 2005), the medial prefrontal cortex (Hartwright et al., 2013; Krause et al., 2012; Völlm et al., 2006), and the posterior superior temporal sulcus (Otsuka et al., 2009; Yang et al., 2015).

Evidence has recently emerged that autistic adults process social information in naturalistic, or ‘real-life’ contexts differently than typically-developing individuals. Several studies have investigated social processing differences between those with and without ASD by examining brain activity in response to watching movies (Bolton et al., 2018; Byrge et al., 2015; Hasson et al., 2009; Salmi et al., 2013). Movie watching mimics real-world experiences by requiring the viewer to integrate perceptual and cognitive systems in order to follow the complexities of the plot. It is known that the brains of healthy individuals become highly synchronized (or correlated) when viewing the same movie (Hasson, Landesman, et al., 2008). This measure of synchronization across different brains is termed *inter-subject correlation* and high levels of synchrony suggest that individuals are experiencing the movie in much the same way. For example, Naci et al. (2014) noted a high degree of synchrony in frontoparietal regions when healthy individuals watched “Bang You’re Dead!” by Alfred Hitchcock and this was shown to relate to how suspenseful and engaging viewers found the movie. The brains of autistic adults have been shown to be less synchronized than those of typically-developing adults during movie watching, and synchrony across individuals tends to be more variable (Bolton et al., 2018; Byrge et al., 2015; Hasson et al., 2009; Salmi et al., 2013). However, this has not been examined in autistic children.

Richardson et al. (2018) have shown that in typically-developing children, those with poorer social cognition have reduced synchrony during movie watching in areas known to be involved with ToM, suggesting that lower synchrony in these areas may also be a feature of autistic children. In the current study, this question was investigated in three groups of children who differed with respect to their social impairment scores and whether or not they had been diagnosed with ASD. Specifically, a data-driven approach was used to examine differences in the degree of inter-subject correlation during movie watching in children aged 7 to 12, who had either been diagnosed with ASD, or did not have ASD but their scores on the Social Responsiveness Scale – revised (SRS-2) indicated social impairments related to autistic traits, or did not have ASD and had typical SRS-2 scores for their age.

On the basis of the existing literature, it was predicted that group differences would emerge in inter-subject correlation within brain networks associated with social cognition. Specifically, it was hypothesized that brain activity within both frontoparietal (Naci et al., 2014), and the ToM networks (Richardson et al., 2018) would be less synchronized in children without ASD with higher social impairment scores than those with lower social impairment scores. Furthermore, it was hypothesized that the brains of children with ASD would be the least synchronized of all, based on their known impairments in many aspects of social cognition.

## Methods

### Dataset

Data was analyzed from the Healthy Brain Network Biobank collected by the Child Mind Institute (described in Alexander et al., 2017), which is an ongoing initiative to collect neuroimaging, medical, and behavioural data on 10,000 participants between the ages of 5 to 21. The Chesapeake Institutional Review Board approved this study. Detailed information on the dataset can be found at http://fcon_1000.projects.nitrc.org/indi/cmi_healthy_brain_network/

### Participants and data acquisition

The Healthy Brain Network Biobank used a community-referred recruitment model to generate a heterogeneous and transdiagnostic sample. Briefly, recruitment involved advertising the study to community members, educators, local care providers, and parents who were on email lists or at events. Potential participants were screened, and were excluded if there were safety concerns, impairments that would interfere with the study procedure (such as being nonverbal or having an IQ of less than 66), and/or medical concerns that could potentially impact brain related findings (for a full description, see Alexander et al., 2017). The study protocol included, where possible, the acquisition of T1 weighted anatomical MRI scans and functional MRI data acquired while the participants watched a ten-minute clip of ‘Despicable Me’ (from 1:02:09 to 1:12:09). All MRI data was collected on a 3T Siemens scanner using a Siemens 32-channel head coil. Functional images were acquired with a gradient-echo planar imaging pulse sequence (TR =800 ms, TE =30 ms, Flip Angle =31 degrees, whole brain coverage 60 slices, resolution 2.4 x 2.4 mm^2^). High-resolution T1-weighted MPRAGE structural images were acquired in 224 sagittal (TR = 2500 ms, TE = 3.15 ms, resolution .8 × .8 mm^2^).

From this database, participants were included in the current analysis if they were between the ages of 7-12 and both anatomical and functional MRI data had been successfully acquired. Everyone included in the current study had written consent obtained from their legal guardians and written assent obtained from the participant. Participants were not excluded based on their handedness. All participants also had scores on the Social Responsiveness Scale Revised (SRS-2), which is a measure of social reciprocity and communication associated with deficits in ASD (Constantino & Gruber, 2012). Specifically, the SRS-2 assesses deficits associated with social awareness, social cognition, social communication, social motivation, and restrictive interests and repetitive behavior, and is rated by parents or caregivers of the child. A score of 59 or below suggests that the child does not have social impairments associated with ASD. A score above 59 is suggestive of impairments in social functioning.

As part of this study, all participants completed a computerized version of the Schedule for Affective Disorders and Schizophrenia - Children’s version (KSADS) in addition to the social responsiveness scale - revised (SRS-2). The KSADS is a semi-structured diagnostic interview used to assess current and past psychopathology according to the DSM-IV criteria, and is rated by a research clinician or social worker (Alexander et al., 2017; Kaufman et al., 1997). Participants who were suspected to have ASD were then assessed in person by a clinician. These participants were assessed using the Autism Diagnostic Observation Schedule – 2^nd^ edition (Lord, Rutter, DiLavore, Risi, Gotham & Bishop, 2012) and the Autism Diagnostic Interview – Revised (Rutter, Le Couteur & Lord, 2003) and those who met the relevant criteria were diagnosed with ASD.

Participants were divided into three groups: The “Low SRS-2 score” (L-SRS) group included those who had an SRS-2 score ≤ 59; the “High SRS-2 Score (H-SRS)” group included participants who had an SRS-2 score of ≥ 60 (the ASD screener cut-off), but were not diagnosed with ASD; and the Autism Spectrum Group (ASD) included participants who were diagnosed with ASD by a clinician as part of the HBN protocol (for details, see Table 1).

**Table 1.** Participant demographics. Means, standard deviations, and ranges of the ages, SRS-2 scores, and the WISC full scale IQ, as well as the number of females and males (F/M) are displayed for each group in the full and matched sample. The full sample of participants was used to create the matched groups. The matched sample was then used for all group comparisons (i.e. the whole brain analysis, the network of interest analysis, and the percentage of synchronized cortex). Only the pairwise cluster-based analysis used the full sample of participants.

| Measure | Group |  |  | Test of group differences |
| --- | --- | --- | --- | --- |
|  | L-SRS | H-SRS | ASD |  |
| <u>Full Sample:</u> |  |  |  |  |
| N | 64 | 34 | 28 |  |
| Age | 9.9 (1.7)<br>7.1 to 12.9 | 9.2 (1.6)<br>7.1 to 11.8 | 9.4 (1.5)<br>7.1 to 12.5 | $F_{(2,123)} = 1.99$,<br>p = .141 |
| Sex (F/M) | 27/37 | 13/21 | 2/26 | $X^2_{(2)} = 11.27$,<br>p = .003 |
| SRS-2 scores | 49.6(4.8)*<br>39 to 59 | 67.5 (6.0)*<br>60 to 84 | 76.6 (10.5)*<br>62 to 90 | $F_{(2, 123)} = 180.98$,<br>p < .001 |
| WISC full scale IQ | 103 (17)<br>61 to 143 | 98 (16)<br>53 to 135 | 93 (18)<br>56 to 129 | $F_{(2,122)} = 3.77$,<br>p = .026 |
| <u>Matched Sample:</u> |  |  |  |  |
| N | 28 | 28 | 28 |  |
| Mean Age | 9.4(1.5)<br>7.0 to 12.6 | 9.6 (1.6)<br>7.2 to 11.8 | 9.4 (1.5)<br>7.1 to 12.5 | $F_{(2,81)} = .155$,<br>p = .857 |
| Sex (F/M) | 3/25 | 7/21 | 2/26 | $X^2_{(2)} = 4.08$,<br>p = .129 |
| SRS-2 | 48.6 (4.8)*<br>41 to 58 | 67.2 (5.7)*<br>60 to 84 | 76.6 (10.5)*<br>62 to 90 | $F_{(2,81)} = 101.79$,<br>p < .001 |
| WISC full scale IQ | 103 (18)<br>61 to 141 | 96 (15)<br>53 to 121 | 93 (18)<br>56 to 129 | $F_{(2,80)} = 2.71$,<br>p = .073 |

Because the groups differed with respect to sample size, age and sex, the L-SRS and H-SRS groups were resampled to produce three demographically matched sub-groups. Specifically, for each participant in the ASD group, an L-SRS and an H-SRS individual who had the same sex and was closest in age (to the month) were selected for inclusion where possible (see Table 1). This resulted in three groups of 28 participants, ensuring sufficient power for acquiring reliable inter-subject correlation results (Pajula & Tohka, 2016). The matched sample was used to statistically compare the groups in the whole brain and network of interest analyses. All but one of the participants in the High SRS-2 group were assessed in person by a clinician. All but one participant also completed the Weschler Intelligence Scale for Children (WISC; Wechsler, 2014).

### MRI pre-processing

For the current study, the MRI data were preprocessed and analyzed using the Automatic Analysis (AA) toolbox (Cusack et al., 2015), SPM8, and in-house MATLAB scripts. Pre-processing of functional data included motion correction (using six motion parameters: left/right, anterior/posterior, superior/inferior, chin up/down, top of head left/right, nose left/right), functional and structural scans were co-registered and normalized to the Montreal Neurological Institute (MNI) template. Functional data were then spatially smoothed using a Gaussian filter (8 mm kernel), and low-frequency noise (e.g., drift) was removed by high-pass filtering with a threshold of 1/128 Hz. The data was denoised using Bandpass filter regressors, with cerebrospinal fluid, white matter signals, motion parameters, their lag-3 2^nd^-order volterra expansion (Friston et al., 2000), and “spikes” (based on mean signal variance across volumes) as nuisance regressors.

### Exploratory whole brain synchronization

To determine the degree of synchronization separately for each group, the degree of inter-subject correlation across the whole brain was calculated using a leave-one-out approach using the matched sample. That is, the pre-processed time course of every voxel was correlated (Pearson and then Fisher z-transformed) between each participant and the mean time course of every voxel from the rest of the group (N-1). A one-sample t-test was calculated on the resulting individual brain-wide correlation values. Multiple comparisons were corrected with a false discovery rate (FDR) of .05 to generate group maps of significantly correlated voxels. To identify where in the brain inter-subject correlation differences existed between the three groups, t-tests were performed on the correlation values at each voxel derived for all of the individuals within each group. Multiple comparisons were corrected with an FDR of .05.

### Network of interest inter-subject correlation

The degree of synchronization within eight previously defined functional networks was calculated. To address our specific hypotheses, a map for the ToM network was used (Dufour et al., 2013) as well as the frontoparietal network from the Yeo et al., (2011) parcellation. Six additional networks (Visual, Dorsal Attention, Ventral Attention, Somatomotor, Limbic, Default Mode Network) from Yeo et al. (2011) were also included in an exploratory analysis to examine potential differences in other areas of the brain. The 8 network parcellations are displayed in Figure 1. Similar to the whole brain inter-subject correlation analysis, the intra-group inter-subject correlation for each of these eight networks was calculated using a leave-one-out approach using the matched sample. Specifically, the time course of each network (based on the average time course of each voxel within the network) for each participant was correlated with the average time course of each network for the remaining participants in the group, minus that participant (N-1). Finally, we used a general linear model to determine if group membership was a significant predictor of intra-group synchronization across the 8 networks. The model included inter-subject correlation values as the predicted variable and group as the predictor variable. This was done separately for each network. The networks that showed a significant effect of group were followed up with Welch t-tests (all results were FDR corrected to .05).

**Figure 1.**
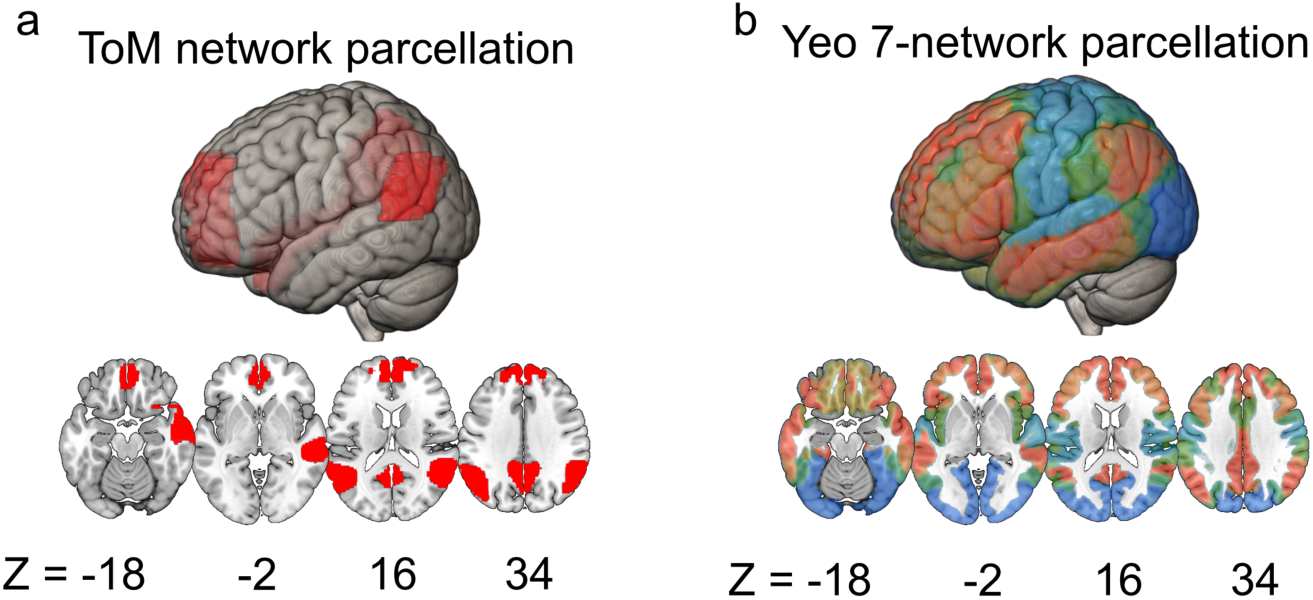
Network of interests used to parcellate the brain. a) ToM network parcellation. The ToM network (displayed in red) included regions in the dorsal, ventral, and lateral medial prefrontal cortex, bilateral temporal parietal junction, precuneus, and right superior temporal sulcus (Dufour et al., 2013). b) The seven network parcellation from Yeo et al. (2011) contained 1) the frontoparietal network (displayed in orange) which includes the lateral prefrontal cortex, medial cingulate, intraparietal sulcus, and inferior temporal gyrus, 2) the visual network (displayed in dark blue) which encompasses the visual cortex, 3) the somatomotor network (displayed in light blue) which includes the motor cortex, premotor cortex, and postcentral gyrus, 4) the dorsal attention network (displayed in dark green) which includes the frontal eye fields, precentral ventral frontal cortex, middle temporal area, and intraparietal sulcus, 5) the ventral attention network (displayed in light green) which includes the dorsal anterior prefrontal cortex, and anterior and posterior cingulate, 6) the limbic network (displayed in mustard) which includes the temporal pole and orbital frontal cortex, 7) the default mode network (displayed in red) which includes the dorsal medial prefrontal cortex, temporal parietal junction, postcentral gyrus, precuneus, the superior temporal sulcus, the posterior cingulate cortex, and retrosplenial cortex.

To better understand the results from the intra-group analysis, the degree of inter-group inter-subject correlation was then calculated, by taking the mean time course for each individual in one group and correlating it with the mean of the two other groups. This generated a correlation value that reflected how similar each participant’s time course was to the two other groups. For instance, we calculated how correlated each ASD participant was to the mean of the other two groups. Finally, we calculated three separate general linear models to determine if participants correlated significantly more with their own group than the mean of the other two groups. The networks that showed a significant effect of group were followed up with Welch t-tests (all results were FDR corrected to .05).

### Percent synchronization across the cortex

The percent of significant voxels across the cortex was calculated, for descriptive purposes, to quantify the number of synchronized voxels common across all individuals in each of the three matched sample groups. To calculate the total percentage of cortex that was synchronized, the number of voxels that were significant per group were divided by the total number of voxels in the brain. To calculate the total percentage of each network that was synchronized, the number of voxels that were significant per group were divided by the total number of voxels in the network of interest.

### Cluster-based inter-subject correlation analysis

To explore the relationship between SRS-2 scores (as a continuous variable) and neural synchrony, pairwise correlations were calculated between each participant and that of every other participant in the ToM and frontoparietal networks. This was done by calculating the mean time course (i.e. the mean activation across each voxel in the network for each time point) in both networks for each participant, and then correlating it with every other participant’s mean time course. Because SRS-2 scores were skewed (upwards) in the ASD and H-SRS groups, this analysis included all participants (N = 126), rather than the smaller matched groups. These pairwise correlations were then plotted in a matrix by ranking each participant by their SRS-2 score (from low to high) for descriptive purposes. Finally, a clustering analysis was conducted to determine whether groups of participants could be identified based solely on their neural synchronization, rather than group membership or SRS-2 scores. To do this, a k-means clustering algorithm was used to group together participants using the time series of neural activity in the ToM and frontoparietal networks. The MATLAB evalclusters function was used to identify the optimal number of clusters based on the variance in the data using the Calinski-Harabasz Index computed over 1000 iterations to minimize the fitting parameter. Based on the groupings generated from this cluster analysis, a logistic regression analysis was computed to investigate which factors (SRS-2 total and subscale scores, age, sex, and social impairment group membership) best predicted the cluster-generated groupings.

## Results

There was a total of 267 eligible participants who met the inclusion criteria (see Methods). Of this sample, 141 participants were removed because of excessive motion, defined as large “spikes”, or significant fluctuations in signal intensity (greater than 3 standard deviations of the mean), in at least 25% of the data.

There was a significant difference between the three groups in terms of SRS-2 scores (F_(2,81)_ = 101.76, p < .001) and post-hoc t-tests showed that the H-SRS group had significantly higher scores than the L-SRS group (t_(51.9)_ = 12.96, p < .001) and had significantly lower scores than the ASD group (t_(42.54)_ = 4.12, p < .001). There were no significant differences between the groups on the WISC full scale IQ scores (F_(2,80)_ = 2.71, *p* = .073), or any of the WISC subscales except for working memory; (F_(2,80)_ = 3.29, *p* = .042). The ASD group had significantly lower working memory scores compared to the L-SRS group (t_(52.10)_ = 2.35, *p* = .023) but not the H-SRS group (t_(50.24)_ = 1.05, *p* = .30).

Differences in correlated motion within each group were examined, in order to ensure that this did not inflate the inter-subject correlation results. Correlated motion was calculated separately for each group, by taking each participant’s 6 motion parameters for each frame and correlating the time course with that of the mean of the rest of the group (N-1). No significant differences were found between the groups in their degree of correlated motion (F_(2,81)_ = .181, p = .835).

### Exploratory whole brain synchronization

Whole brain synchronization was characterized in the three groups. All groups showed significant synchronization in the auditory and visual areas (Figure 2a). In fact, synchronization in these areas was stronger than in any other brain areas, replicating previous inter-subject correlation findings during movie watching (Hasson et al., 2008). The H-SRS and L-SRS groups also showed significant inter-subject correlation in areas associated with ToM and executive processing, including parts of the right and left temporal parietal junction, the precuneus, the intraparietal sulcus, the superior parietal lobe, and portions of the medial and lateral prefrontal cortex. In contrast, the ASD group had very little significant inter-subject correlation outside of visual and auditory areas (see Figure 2a, bottom row).

**Figure 2.**
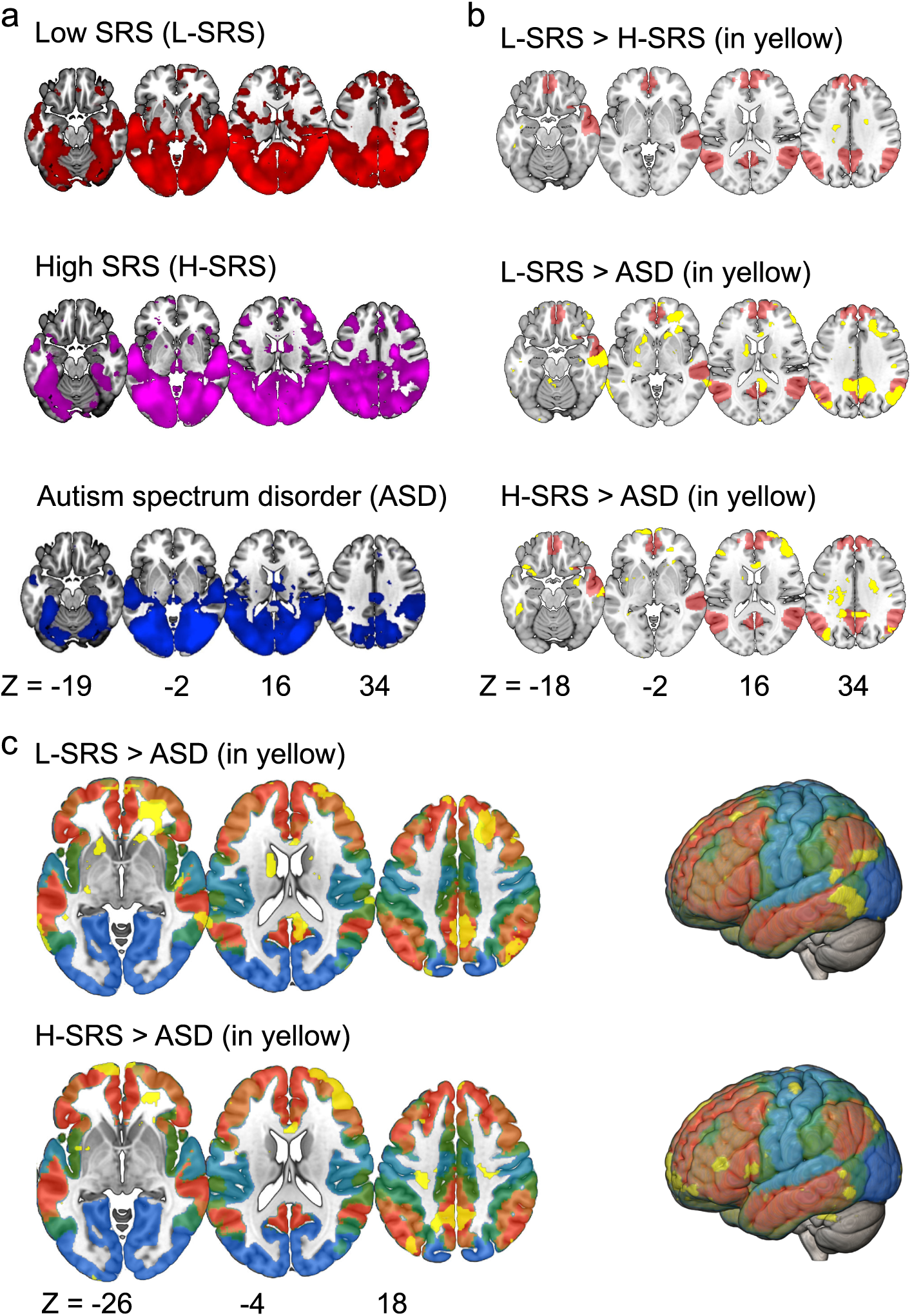
Exploratory whole brain inter-subject correlation analysis. **a)** Voxels displayed in red showed significant inter-subject correlation during movie watching in the L-SRS group. Voxels displayed in violet showed significant inter-subject correlation in the H-SRS group. Voxels displayed in blue showed significant inter-subject correlation in the ASD group. All p values were FDR corrected to an alpha of .05. **b)** Whole brain contrasts were calculated by conducting one-tailed t-tests on the inter-subject correlation values between each group (p values corrected to an FDR of .05). Voxels displayed in yellow showed significantly greater inter-subject correlation values based on this contrast, voxels displayed in red show the ToM network parcellation. c) Voxels displayed in yellow showed significantly greater inter-subject correlation values based on the same contrast displayed in b, overlaid on top of the Yeo et al. (2011) 7-network parcellation. (Frontoparietal = orange, Visual = dark blue, Somatomotor = light blue, Dorsal attention = dark green, Ventral attention = light green, Limbic = mustard, Default mode = red).

Next, whole brain contrasts were conducted (Figure 2b) to examine whether the magnitude of synchronization differed between the three groups. When the L-SRS group was contrasted to the H-SRS group, only tiny areas of difference were observed after multiple comparisons corrections, in the inferior temporal gyrus (MNI coordinates _X, Y, Z_ = -46, -37, -17, *t*_(54)_ = 4.61, p_corrected_ = .030), and white matter (see Figure 2b, top row). The L-SRS group showed significantly greater inter-subject correlation than the ASD group (Figure 2b, middle row) in the bilateral temporal parietal junction (MNI coordinates (left) = -57, -61, 30, *t*_(54)_ = 3.97, p_corrected_ = .011, MNI coordinates (right) = 49, -67, 31, *t*_(54)_ = 5.07, p_corrected_ = .002), precuneus (MNI coordinates = 4, -51, 41, *t*_(54)_ = 4.14, p_corrected_ = .009), right superior temporal sulcus (MNI coordinates =60, -11, -16, *t*_(54)_ = 4.47, p_corrected_ = .005), right hippocampus (MNI coordinates = 32, -14, -19, *t*_(54)_ = 3.52, p_corrected_ = .026), and in regions of the lateral (MNI coordinates = 39, 54, -9, *t*_(54)_ = 3.61, p_corrected_ = .022), and the right medial prefrontal cortex (MNI coordinates = 22, 42, 37, *t*_(54)_ = 3.89, p_corrected_ = .014). The H-SRS group had significantly greater synchronization than the ASD group in the precuneus (MNI coordinates = -3, -55, 45, *t*_(54)_ = 4.00, p_corrected_ = .017), right hippocampus (MNI coordinates = 28, -5, -21, *t*_(54)_ = 3.37, p_corrected_ = .043), and in regions of the lateral (MNI coordinates = 46, 44, 12, *t*_(54)_ = 6.30, p_corrected_ < .001), and medial prefrontal cortex (MNI coordinates = -5, 65, -7, *t*_(54)_ = 4.27, p_corrected_ = .012) (Figure 2b, bottom row).

### Network based synchronization

Group differences in the magnitude of intra-group synchronization revealed a main effect of group in the ToM (F _(2,81)_ = 4.94, *p* = .009) and the limbic (F _(2,81)_ = 3.93, p = .023) networks (Figure 3), but not in any of the others examined, including the frontoparietal network (F _(2,81)_ = 2.02, *p* = .140, Cohen’s d ranged from .037 to .476). Post-hoc analyses of neural synchronization revealed that the ASD group had significantly lower inter-subject correlation values compared to the L-SRS group within the ToM (t _(50.11)_ = 3.50, *p*_corrected_ = .006, Cohen’s d = .934) and limbic networks (t _(50.00)_ = 2.48, *p*_corrected_ = .044, Cohen’s d = .664). They also had significantly lower inter-subject correlation values compared to the H-SRS group in the limbic network (t _(50.21)_ = 2.18, *p*_corrected_ = .044, Cohen’s d = .631), although differences in inter-subject correlation just failed to meet the corrected alpha level in the ToM network (t _(52.33)_ = 2.36, *p*_corrected_ = .0504, Cohen’s d = .584). Moreover, no significant differences in inter-subject correlation were observed between the L-SRS and H-SRS groups within the ToM (t _(45.72)_ = .488, *p*_corrected_ =.628, Cohen’s d = .130) or limbic networks (t _(45.21)_ = .417, *p*_corrected_ =.628, Cohen’s d = .111).

**Figure 3.**
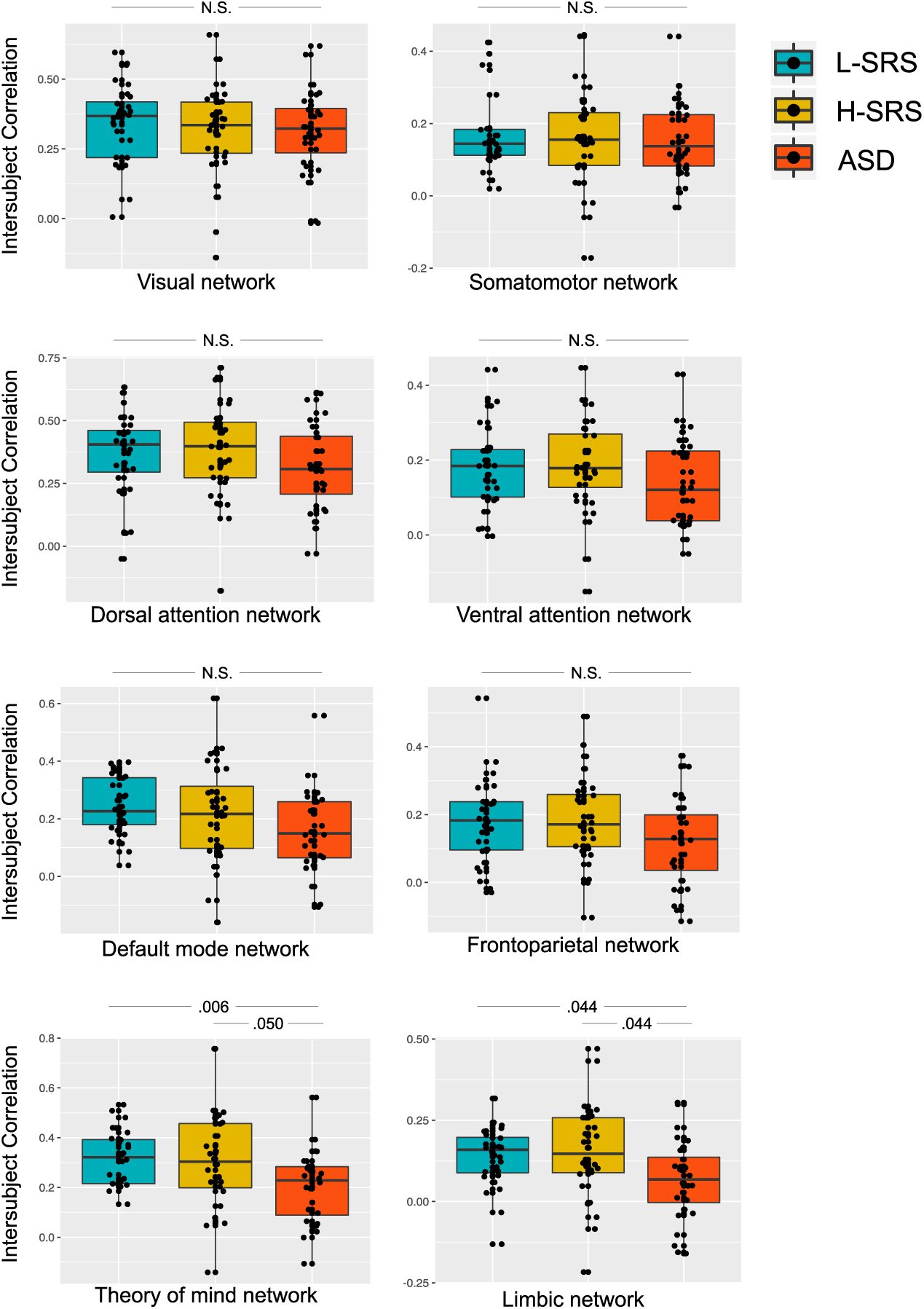
Intra-group network of interest analysis. Mean inter-subject correlation, based on the leave one out correlation analysis conducted separately for each group, is displayed as dots for each participant in the eight networks. Boxplots indicate the median inter-subject correlation value and interquartile range for each group (blue = L-SRS, yellow = H-SRS, red = ASD). The ASD group had significantly lower inter-subject correlation in the limbic and ToM networks compared to the L-SRS group. The ASD group also had significantly lower inter-subject correlation in the limbic network compared to the H-SRS group, while in the ToM network this difference narrowly missed statistical significance (corrected p value = .0504). The groups did not differ significantly in any of the six other networks.

An inter-group inter-subject correlation network analysis was performed to investigate whether individuals in one group had significantly greater neural synchronization with their own group than that of the other two groups. The results revealed that the degree of inter-subject correlation was not significantly different between any of the groups in any of the examined networks, including the frontoparietal and ToM networks.

### Percent synchronization across the cortex

When looking at the percentage of synchronized voxels across the whole brain, the ASD group had nearly one-third less (38%) than the L-SRS (56%) and H-SRS (52%) groups (see Figure 4). The percentage of significant voxels in each of the eight networks of interest was also calculated (see Figure 4). The difference in percentage across the whole brain between the groups was not accounted for by less synchronization in any one network; rather, the ASD group had fewer synchronized voxels in every network, including in the ToM and frontoparietal networks.

**Figure 4.**
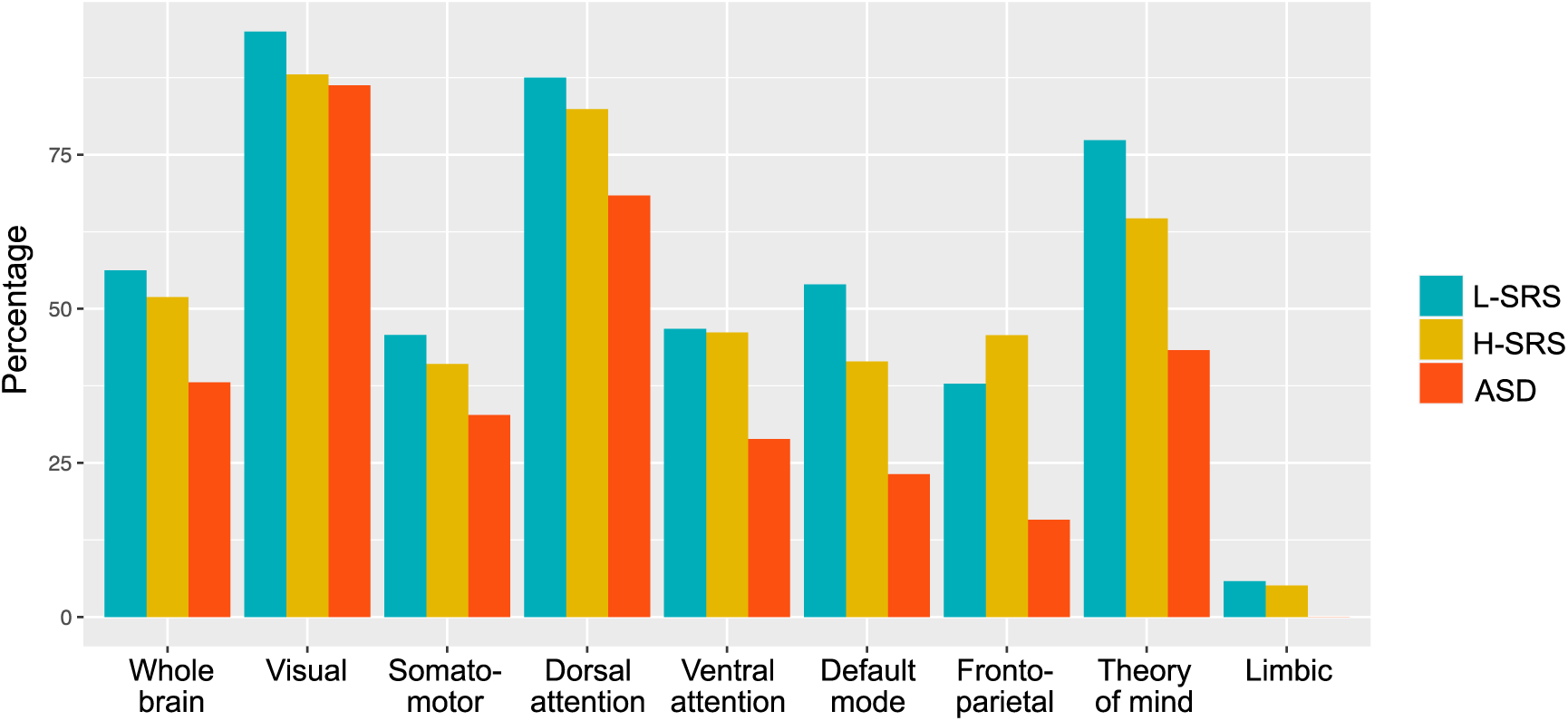
Percentage of correlated voxels. The percent of significant voxels across the cortex was calculated, for descriptive purposes, to quantify the number of synchronized voxels common across all individuals in each of the three matched sample groups. This was calculated by dividing the number of voxels with significant inter-subject correlation by the total number of voxels in the whole brain or network for each group separately (blue = L-SRS, yellow = H-SRS, red = ASD).

### Cluster-based inter-subject correlation analysis

To explore whether SRS-2 scores predicted inter-subject correlation values when used as a continuous measure (instead of a categorical variable), pairwise inter-subject correlations were calculated between each participant (N=126) in the frontoparietal and ToM networks. The entire sample was used so that the SRS-2 scores were normally distributed and to increase statistical power. Pairwise correlations were conducted to reduce any influence the groupings may have had on the mean time course originally used to calculate inter-subject correlation. For instance, if those with low SRS-2 scores and those with high SRS-2 scores both correlated with their own group to a similar degree, but the pattern of activations was different, using these groupings would obfuscate any differences. For descriptive purposes, the matrix of pairwise correlation values was plotted by ranking each participant by their SRS-2 score, from low to high (see Figure 5a). A k-means clustering analysis was conducted on the pairwise correlations to explore potential factors that predicted groups of participants who have the most similar degree of synchrony in these two networks. The best fit was achieved by dividing the data into two clusters in both the frontoparietal and ToM networks; cluster 1 included individuals with similar neural responses to the movie (large positive correlations) and cluster 2 included individuals with unrelated neural responses to the movie (Figure b). Moreover, there was also a large overlap between the participants who were in cluster 1 in the ToM and frontoparietal networks. Specifically, of the 58 participants who had high similarity in the ToM network (cluster 1), 45 of them also had high similarity in the frontoparietal network.

**Figure 5.**
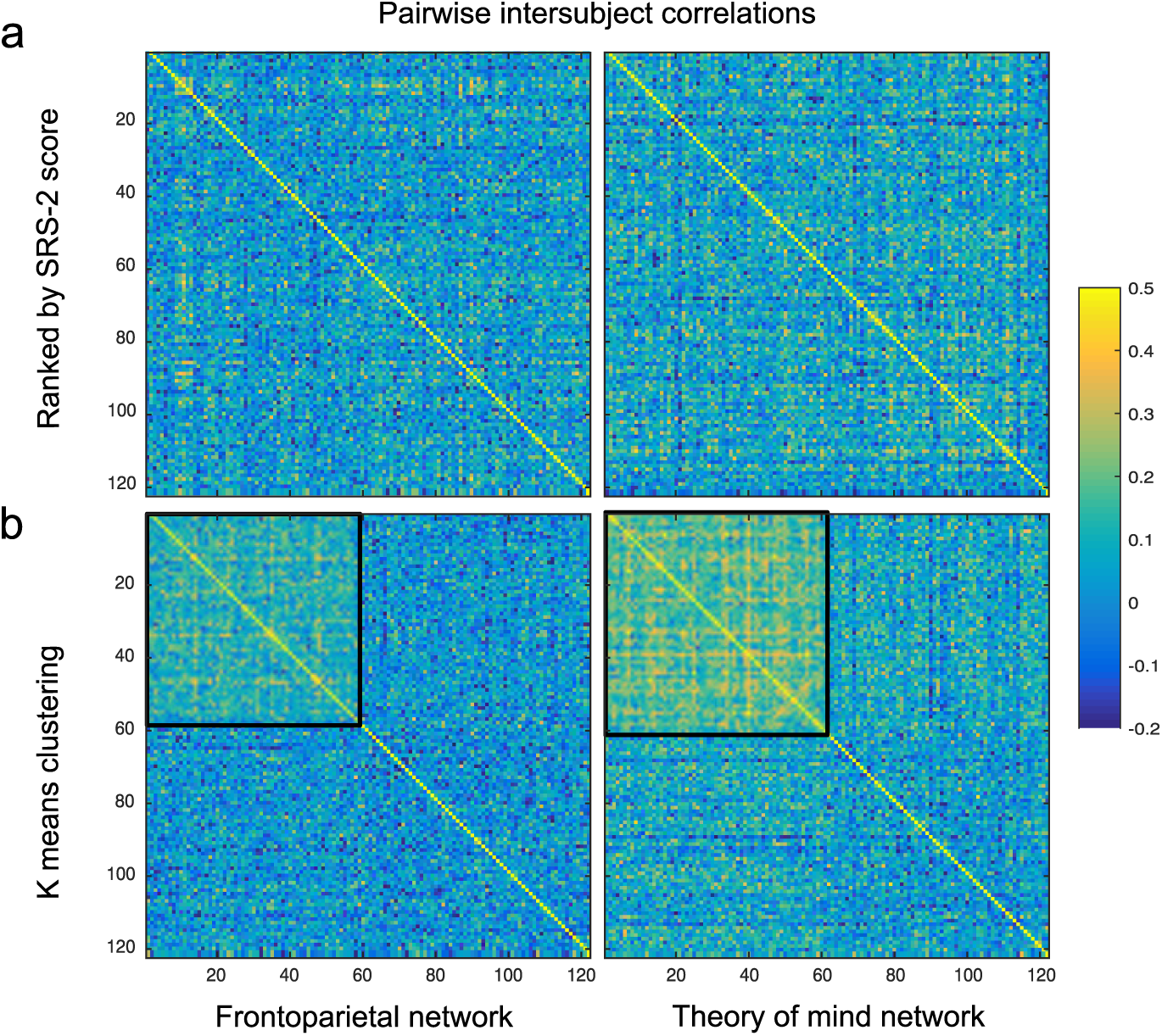
Pairwise inter-subject correlations. Yellow squares indicate a higher positive correlation (i.e. high similarity in time series), blue squares indicated a low or negative correlation (i.e. low similarity in time series). **a)** Pairwise correlations in time series in the frontoparietal and ToM networks between each pair of participants are ordered by SRS-2 scores (from low to high). **b)** Pairwise correlations in time series are ordered based on the K-means analysis in the frontoparietal and ToM networks. Black boxes show cluster 1 (the high similarity group) for each network.

Logistic regression was run to determine whether the probability of being in cluster 1 versus cluster 2 could be predicted by age, sex, full scale IQ, SRS-2 total and subscales, or group membership (i.e., L-SRS, H-SRS and ASD). None of these factors significantly predicted cluster membership in the frontoparietal network. However, in the ToM network, group membership significantly predicted cluster membership. Cluster 1 comprised 35 participants (60%) in the L-SRS group, 17 individuals (29%) from the H-SRS groups, and 6 individuals (10%) who were diagnosed with ASD. In contrast, cluster 2 consisted of 29 individuals (45%) from the L-SRS group, 17 individuals (20%) from the H-SRS group, and 22 individuals (35%) diagnosed with ASD. There were significantly more participants from the ASD group in cluster 2 than in cluster 1 in the ToM network (X^2^_(1)_ = 7.5, p = .006), while there was no significant difference in the number of H-SRS participants between the two clusters (X^2^_(1)_ = .11, p = .73), and although there were more L-SRS participants in cluster 1, this difference did not reach significance (X^2^_(1)_ = 3.25, p = .072).

## Discussion

In the current study, a group of ASD participants had significantly less neural synchronization when watching a movie compared to the L-SRS and H-SRS groups across the whole brain, including the ToM and limbic networks, as well as the lateral and medial prefrontal cortex. These regions have been shown previously to be associated with elements of ‘plot following’ during movie watching (Hasson, Furman, et al., 2008; Hasson, Landesman, et al., 2008; Naci et al., 2014; Nguyen et al., 2019), suggesting that the children in the ASD group were experiencing the movie qualitatively differently than the participants in the other two groups. These results, in particular the fact that the ToM network was less synchronized in the ASD group, are intriguing given that regions within this network are associated with social cognition (Dufour et al., 2013; Mills et al., 2014; Richardson et al., 2018; Rilling et al., 2004), which is known to be affected in ASD (Hamilton et al., 2009; Pedreño et al., 2017; Spencer et al., 2011). While aspects of social cognition are usually discussed in the context of inter-personal relationships, they are also essential components of movie-watching, allowing one to become immersed in the plot by taking the perspective of the characters appropriately, understanding their motives, and following their verbal and nonverbal communication cues. Yeshurun et al. (2017) have reported previously that manipulating an individual’s understanding of a plot reduces neural synchrony in ToM regions, including the precuneus, temporal parietal junction, and medial prefrontal cortex. Thus, these findings support the idea that autistic children process social stimuli in a distinct way, as they have different neural responses in the ToM network during a movie, when compared to children without ASD.

It is also interesting that participants in the ASD group had significantly less synchrony in the lateral prefrontal cortex, a region within the frontoparietal network, when compared to those in the other two groups. Understanding a complex narrative (such as a movie’s plot) requires a viewer to remember previous events, pay attention to what is currently happening, make predictions about the future consequences of current events, and integrate this information over time, all of which depends on frontoparietal executive processing (Naci et al., 2014). In previous studies, reduced synchrony in this network has been associated with ‘losing the plot’ during deep sedation (Naci et al., 2018), and in patients with severe brain damage (Naci et al., 2014). Thus, this decrease in inter-subject correlation in the lateral prefrontal cortex may suggest that participants in the ASD group are also failing to grasp elements of the plot in the way that the other participants do.

Despite finding that inter-subject correlation was reduced in prefrontal regions using a whole brain analysis, no differences in the degree of inter-subject correlation were found in the frontoparietal network when a network of interest analysis was used. One potential reason is that the parcellation used for the frontoparietal network was based on adult data and may not accurately capture this network in children. Previous work has shown that the frontoparietal network continues to develop into early adulthood (Baum et al., 2017; Peters et al., 2016), and so the parcellation masks from Yeo et al. (2011) may have led us to average neural activity from regions that are not yet fully integrated in children.

While not part of our hypotheses, it is interesting that the ASD group showed less inter-subject correlation in the right hippocampus in the whole brain analysis as well as in the limbic network, when examined using the parcellation by Yeo et al. (2011). Similar findings have been reported in autistic adults watching movies (Byrge et al., 2015). Moreover, Chen et al. (2017) found that, in healthy adults, the degree of inter-subject correlation within the hippocampus during movie watching predicted events that were later recalled, although this has not been examined during development. Nevertheless, long-term memory deficits have been reported in ASD; specifically, autistic individuals perform worse on episodic, but not semantic, memory tasks (Crane & Goddard, 2008; Lind, 2010)

Contrary to our hypothesis, no meaningful differences in neural synchrony were found between the L-SRS and H-SRS groups. This contrasts with the results of Richardson et al., (2018) who found that social cognition in typically-developing children was related to the degree of inter-subject correlation within the ToM network during movie-watching. One potential reason for this difference is that Richardson et al., (2018) calculated inter-subject correlation based on how similar each child’s time course was to a group of adults watching the same movie, whereas in the current study, inter-subject correlation was calculated by correlating each participant’s time course to the mean of their own group. Moreover, the measure of social cognition used by Richardson et al. (2018) focused specifically on comprehension of a social narrative, which has many things in common with how people follow the plot of a movie. It is perhaps not surprising then, that the two things correlated. In the current study, a measure of social impairment was used – the SRS-2, which measures an individual’s motivation to engage in social interactions, their use of social communication, and their ability to understand social cues (Constantino & Gruber, 2012). Thus, while the H-SRS and L-SRS groups differed in terms of their social impairments as measured by the SRS-2 scale, these mechanisms may be unrelated, or only moderately related, to those that are involved in plot following. Moreover, it is also possible that creating categorical groups based on the SRS-2 scores may have obscured subtle differences in individuals with differing levels of social impairment. To investigate this possibility, the exploratory pairwise correlation analysis was conducted, which found that SRS-2 scores as a continuous measure did not predict whether participants had similar patterns of neural activity in the ToM or frontoparietal networks. Taken together, these results suggest that it is only when social impairment is in the clinical range, as is seen in ASD, that differences in conscious processing of naturalistic stimuli emerge.

As a group, autistic participants had less inter-subject correlation compared to those without ASD, but these differences did not apply uniformly to each individual. The clustering analysis indicated that the majority of ASD participants had low similarity in their time courses compared to all other participants. However, six out of 28 of those diagnosed with ASD clustered with the ‘high similarity’ group (comprising about 10% of the group) according to their synchronization in the ToM network. Using a similar clustering analysis, Byrge et al., (2015) found that in a sample of 17 high functioning autistic adults, five showed idiosyncratic patterns of inter-subject correlation compared to typically-developing individuals, while the other 12 clustered with the control group. Moreover, they found that these five individuals were significantly worse than the control group and the other 12 ASD participants, when asked to explain elements of a movie plot. Together, these findings suggest that lower synchronization during movie-watching may be common, but not a uniform characteristic of either autistic children or adults. Indeed, heterogeneity in clinical features, cognitive profiles, and differing genetic and environmental risk factors has plagued research in ASD (Betancur, 2011; Jeste & Geschwind, 2014; Lenroot & Yeung, 2013). For example, within the neuroimaging literature, some studies have reported underconnectivity across the brains of autistic individuals (Cherkassky et al., 2006; Di Martino et al., 2014; von dem Hagen et al., 2013), while others find hyperconnectivity (Supekar et al., 2013; Uddin et al., 2010, 2013).

Finally, it is important to keep in mind the exploratory nature of the current study when interpreting these findings. This is a step towards a better understanding of how children with and without ASD process naturalistic stimuli, but replication and further investigation is needed to better understand the nature of the differences observed. For instance, one potential mechanism underlying our results could be that participants in the ASD group had more variable neural responses to the movie. However, it would be valuable for future studies to directly examine if more variable neural responses to movies are driving reduced neural synchronization in those diagnosed with ASD. Additionally, a major limitation of this study is that no memory test, or measure of how well the movie clip was understood, was collected. A behavioral measure of movie comprehension may help to explain the nature of the neural differences observed in this study. It is possible that individuals were attending to different features of the movie, which has been shown to influence the degree of neural synchrony (Nguyen et al., 2019), although previous work has confirmed that movies similar to ‘Despicable me’ maintain the viewers’ attention (Hasson, Landesman, et al., 2008; Naci et al., 2015). It is also unlikely that participants were asleep during the movie, as most of the visual network was synchronized across the three groups during the movie, which is not observed when individuals are sedated (Naci et al., 2018).

## Conclusion

In sum, the current results suggest that autistic children, as a group, process movies in a unique way compared to those without ASD. Interestingly, a minority of these children had time courses that were highly correlated with a group of children without ASD in the ToM network. Future research should investigate factors that underlie this heterogeneity, as this may be one avenue to better understand how autistic individuals process the world around them.

## Author contributions

Conceptualization, writing–reviewing and editing, K.M.L., R.A.S, A.M.O, B.S.; Methodology, Formal analysis, K.M.L., B.S.; writing–original draft preparation, K.M.L., B.S., A.M.O; funding acquisition, A.M.O., B.S

## Acknowledgements

The authors thank the Child and Mind Institute for designing and collecting data for the Healthy Brain Biobank.

## Funding

This research was funded by the Canada Excellence Research Chairs Program (#215063), the Canadian Institutes of Health Research (#408004), and the Natural Sciences and Engineering Research Council of Canada (#390057).

## Conflict of interest

The authors have no conflicts of interest to declare.

## Abbreviations

ASD: Autism spectrum disorder
fMRI: functional magnetic resonance imaging
L-SRS: High SRS-2 score
SRS-2: Low SRS-2 score
H-SRS: Social responsiveness scale – revised
ToM: Theory of mind
WISC: Weschler Intelligence Scale for Children

## Notes

### Competing Interest Statement

The authors have declared no competing interest.

